# Rif1 functions in a tissue-specific manner to control replication timing through its PP1-binding motif

**DOI:** 10.1101/870451

**Authors:** Robin L. Armstrong, Souradip Das, Christina A. Hill, Robert J. Duronio, Jared T. Nordman

## Abstract

Replication initiation in eukaryotic cells occurs asynchronously throughout S phase, yielding early and late replicating regions of the genome, a process known as replication timing (RT). RT changes during development to ensure accurate genome duplication and maintain genome stability. To understand the relative contributions that cell lineage, cell cycle, and replication initiation regulators have on RT, we utilized the powerful developmental systems available in *Drosophila melanogaster*. We generated and compared RT profiles from mitotic cells of different tissues and from mitotic and endocycling cells of the same tissue. Our results demonstrate that cell lineage has the largest effect on RT, whereas switching from a mitotic to an endoreplicative cell cycle has little to no effect on RT. Additionally, we demonstrate that the RT differences we observed in all cases are largely independent of transcriptional differences. We also employed a genetic approach in these same cell types to understand the relative contribution the eukaryotic RT control factor, Rif1, has on RT control. Our results demonstrate that Rif1 can function in a tissue-specific manner to control RT. Importantly, the Protein Phosphatase 1 (PP1) binding motif of Rif1 is essential for Rif1 to regulate RT. Together, our data support a model in which the RT program is primarily driven by cell lineage and is further refined by Rif1/PP1 to ultimately generate tissue-specific RT programs.

## Introduction

DNA replication initiates from discrete regions of the eukaryotic genome, known as replication domains, in a precise chronological manner during S phase. This temporal order of DNA replication is known as the DNA replication timing (RT) program and is evolutionarily conserved from yeast to humans (Rivera-Mulia and Gilbert 2016). In metazoan species, replication domain sizes range from hundreds of kilobases to megabases, and their RT is correlated with transcriptional activity, chromatin structure, and position within the nucleus (MacAlpine et al. 2004; Schwaiger et al. 2009; Eaton et al. 2011; Rivera-Mulia and Gilbert 2016; Almeida et al. 2018). Furthermore, RT domains are highly correlated with topologically associated domains (TADs), where a near one-to-one correlation has been observed between RT domains and TADs (Pope et al. 2014). While RT is clearly influenced by chromatin structure and nuclear organization, the exact function of RT is not fully understood. Importantly, defects in RT are associated with genome instability, and RT is often altered in cancer cells (Stamatoyannopoulos et al. 2009; Koren et al. 2012; Donley and Thayer 2013). Therefore, understanding the processes and factors that contribute to RT is key to understanding fundamental aspects of eukaryotic DNA replication and genome stability.

Both cellular differentiation and cellular identity influence genome-wide RT, suggesting that the underlying mechanisms regulating RT are plastic during development. Comparison of genome-wide RT between three lines of cultured *Drosophila* cells revealed differences in RT across ~8% of the genome (Lubelsky et al. 2014). More extensive RT profiling using *in vitro* models of cellular differentiation from multiple mammalian cell lineages has revealed ~50% of the genome is subject to cell-type specific RT changes (Hiratani et al. 2008; Hiratani et al. 2010). Furthermore, in mammalian cells, the RT program goes through a global reorganization where many small RT domains consolidate into larger RT domains as cells differentiate from embryonic stem cells to more differentiated cell types (Ryba et al. 2010). It is still unclear, however, whether cell-type specific changes in RT are developmentally programmed directly or whether differential RT is a passive reflection of the changes in chromatin structure and nuclear organization that occur during cellular differentiation.

Multiple *trans*-acting replication factors control RT from yeast to humans. Loading of the MCM replicative helicase during G1 phase of the cell division cycle and helicase activation during S phase are key steps in RT control (Bell and Stillman 1992; MacAlpine et al. 2010; Mantiero et al. 2011; Collart et al. 2013; Miotto et al. 2016). Several factors are limiting for replication initiation (Sld2, Sld3, Dpb11, Dbf4 and Cdc45) and their overexpression disrupts RT in budding yeast and *Xenopus* (Mantiero et al. 2011; Collart et al. 2013). A critical *trans*-acting RT-regulating factor is Rif1 (<u>R</u>ap1-interacting factor <u>1</u>), which controls RT from yeasts to humans (Cornacchia et al. 2012; Hayano et al. 2012; Yamazaki et al. 2012; Peace et al. 2014; Foti et al. 2016). In animals, it is not clear whether the genomic regions that Rif1 targets during differentiation are cell-type specific or whether Rif1 selectively regulates specific regions of the genome regardless of cell type. Although Rif1 is only modestly conserved, all Rif1 orthologs contain a <u>P</u>rotein <u>P</u>hosphatase <u>1</u> (PP1)-interaction motif, suggesting that PP1 recruitment is a critical function of Rif1. Rif1-dependent recruitment of PP1 to chromatin may prevent the Dbf4-dependent kinase (DDK) activation of loaded helicases (Davé et al. 2014; Hiraga et al. 2014; Mattarocci et al. 2014; Hiraga et al. 2017; Sukackaite et al. 2017). How loss of the Rif1-PP1 interaction affects RT genome wide, however, has not been determined.

To better understand the extent to which Rif1 regulates RT in various unperturbed cell types during development, we have measured RT in the *Drosophila* larval wing discs and adult ovarian follicle cells in the presence and absence of Rif1. Here, we identify regions of the genome that change RT as a function of cell lineage and determine Rif1-dependent changes in RT in different tissue types. We found that cell lineage is a major driver of RT and demonstrate that tissue-specific transcription is not a major contributor to tissue-specific RT. Importantly, although RT in a subset of the genome depends on Rif1 similarly in different tissues, Rif1 acts in a tissue-specific manner to control RT. Additionally, the Rif1-PP1 interaction motif is required for Rif1-dependent control of RT, suggesting that PP1 recruitment to replicative helicases is the predominant mechanism Rif1 utilizes for RT control.

## Results

### Cell lineage is a major driver of DNA replication timing

To analyze RT in unperturbed cell types and tissues without the need to immortalize or transform cells, we exploited the well-characterized developmental systems of *Drosophila melanogaster*. To determine how cell lineage affects RT, we generated genome-wide RT profiles from cells of two distinct *D. melanogaster* epithelial tissues: third-instar larval wing imaginal disc cells and follicle cells from female adult ovaries. Cells of the wing disc are derived from the embryonic mesoderm while ovarian follicle cells are derived from the embryonic ectoderm. To generate RT profiles, we used fluorescence-activated cell sorting (FACS) to isolate and subsequently sequence the genomes of S phase nuclei from each tissue and compared these data to those obtained from G1 phase nuclei from wing discs (Figure 1A; (Armstrong et al. 2018)). The premise of this method is that early-replicating DNA sequences are over-represented relative to late-replicating sequences within the S phase population. Therefore, replication timing values can be quantified by determining log_2_ transformed S/G1 read counts across the genome, where larger values indicate earlier replication and smaller values indicate later replication (Figure 1A).

**Figure 1.**
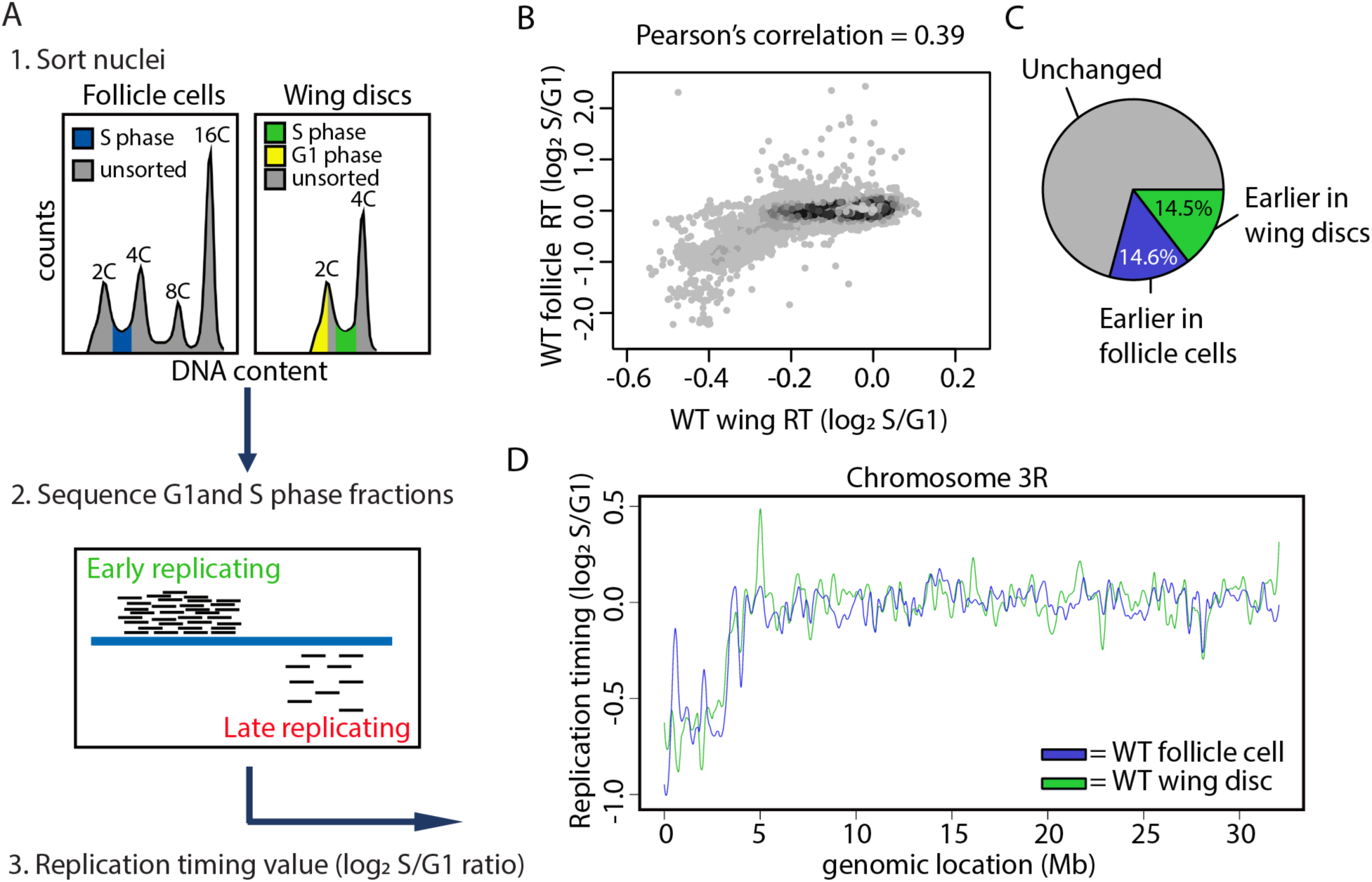
Cell lineage is a major driver of DNA replication timing in *Drosophila*. **A)** Experimental outline: (1) Nuclei were FACS sorted into G1 (yellow) and S (blue or green) populations based on DNA content. (2) DNA was sequenced and mapped back to the dm6 reference genome. More reads map to early than late replicating sequences. (3) S/G1 log_2_ ratio of mapped reads generates replication timing profiles. **B)** Heatscatter plot of wildtype wing disc and wildtype follicle cell S/G1 (log_2_) ratios at all 100kb windows using a 10kb slide across the genome. **C)** Pie chart of all 100kb windows of significantly earlier RT in wildtype wing discs (green), significantly earlier RT in wildtype follicle cells (blue), and unchanged RT (grey) across the major chromosome scaffolds. **D)** LOESS regression lines showing average wildtype wing disc (green) and wildtype follicle cell (blue) S/G1 (log_2_) replication timing values across the chromosome 3R scaffold. See Figure S1 for all other chromosome arms.

To determine how lineage contributes to RT, we generated RT values at 100kb windows tiled at 10kb intervals across the genome for both wing discs and follicle cells and used a stringent significance threshold to identify differential RT between each tissue (Materials and Methods; (Armstrong et al. 2018)). RT profiles generated from individual replicates of wildtype wing discs and follicle cells were strongly correlated (Pearson’s correlations = 0.95 and 0.95, respectively; Figure S1A), whereas RT values between the two lineages were significantly more divergent (Pearson’s correlation = 0.39; Figure 1B). While ~70% of the genome has similar RT between the two tissues, ~29% of the genome displays tissue-specific RT where 14.6% of windows replicate earlier in follicle cells and 14.5% of windows replicate earlier in wing discs (Figure 1C,D; Figure S1B; Table S1). Gene ontology analysis of genes located within tissue-specific RT domains did not reveal a significant enrichment of genes associated with a specific biological process. Furthermore, differential RT between wing discs and follicle cells did not preferentially affect any one chromatin state (Kharchenko et al. 2011), and replication domain sizes were highly similar between the two tissues (Figure S1C,D). These data demonstrate that cell lineage is a key contributor to replication timing control in *Drosophila* similar to what has been previously observed in mammalian cell culture systems (Hiratani et al. 2008; Ryba et al. 2010; Rivera-Mulia et al. 2015).

### Cell type-specific transcription does not drive changes in RT

Transcriptional activity is highly correlated with RT, with early replicating regions of the genome associated with active transcription and late replicating regions associated with transcriptional repression (MacAlpine et al. 2004; Liu et al. 2012; Lubelsky et al. 2014; Rivera-Mulia and Gilbert 2016). Therefore, we determined if differences in transcriptional activity are correlated with differential RT. We generated transcriptomes from wildtype wing disc cells and follicle cells by total RNA-seq and identified differentially expressed transcripts between each tissue type. Individual biological replicates were highly correlated (Figure S2; Pearson’s correlation coefficients > 0.95) and we were able to identify tissue-specific gene expression including *wingless* (*wg*) expression in wing discs and *chorion protein* (*cp*) expression in follicle cells (Figure S3A). We observed 3,994 differentially expressed transcripts (p < 0.01; edgeR) between the two tissues (Figure 2A), with elevated expression of 2,651 transcripts in wing discs and 1,343 transcripts in follicle cells (Figure 2A).

**Figure 2.**
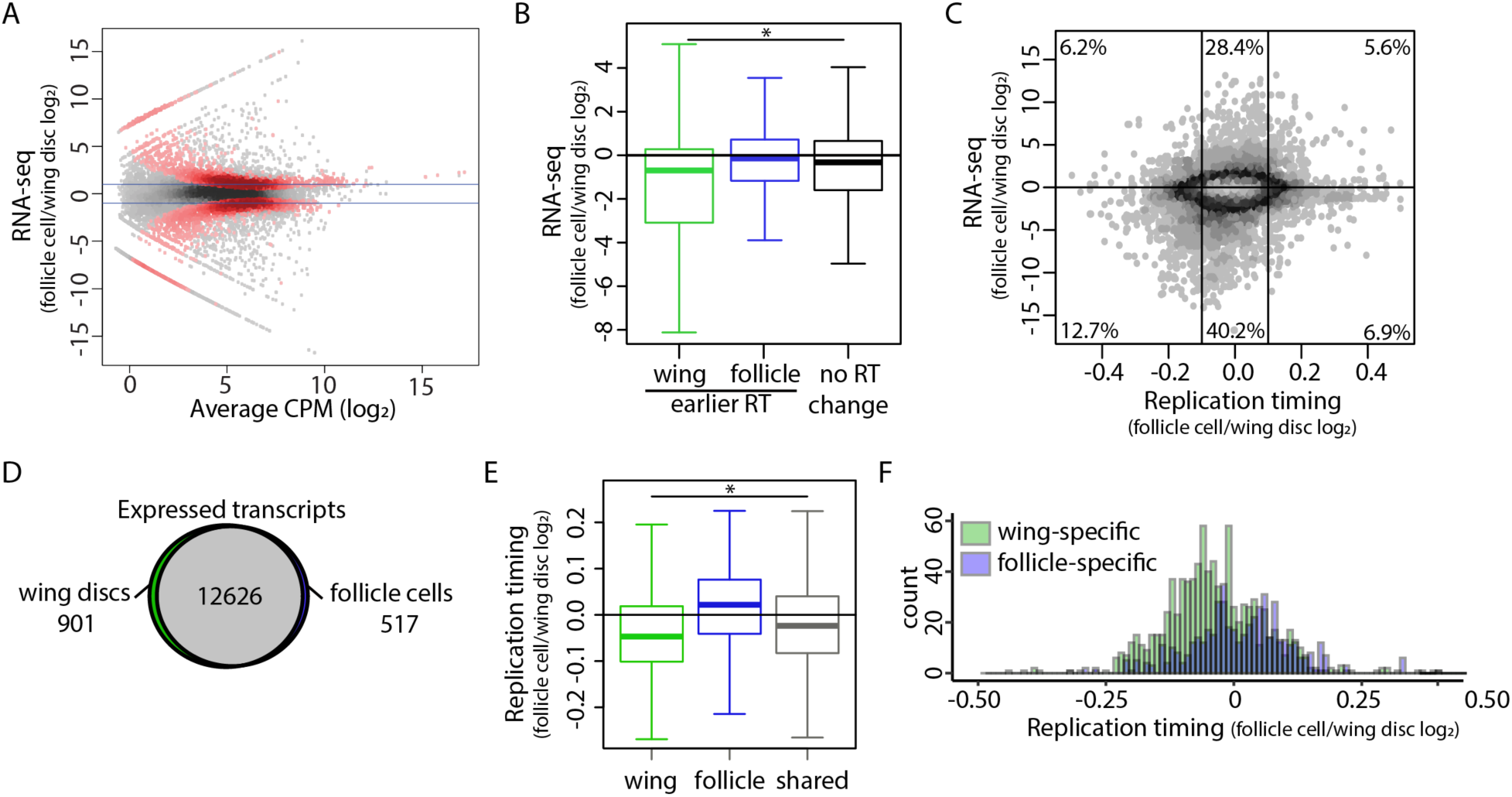
Tissue-specific transcription does not drive changes in RT. **A)** Heatscatter plot of the wildtype follicle cell/wildtype wing disc ratio of total RNA-seq signal. Statistically different transcripts between wildtype follicle cells and wildtype wing discs are indicated in red (p < 0.01; edgeR). Blue lines indicate a log_2_ fold change of 1 and −1. **B)** The average log_2_ fold change of all transcripts within each 10kb window of earlier RT in wildtype wing discs (green), earlier RT in wildtype follicle cells (blue), and unchanged RT (grey). Only windows containing at least one transcript are shown. (p < 0.0001; One way ANOVA). **C)** Heatscatter plot of the wildtype follicle cell/wildtype wing disc RT values (S/G1 (log_2_)) versus the wildtype follicle cell/wildtype wing disc ratio of normalized RNA-seq signal at all 10kb windows across the major chromosome scaffolds. The average log_2_ fold change of all transcripts within each 10kb window is plotted, and only windows containing at least one transcript are shown. Percentages represent the number of windows within each region (vertical lines at −0.1 and 0.1 represent log_2_ fold change cutoffs for RT statistical significance). **D)** Venn diagram comparing expressed transcripts (TPM > 0) between wildtype wing discs and wild type follicle cells. Wing-specific (green), follicle-specific (blue) and shared (grey) transcripts are indicated. **E)** Log_2_ fold change of RT values between wildtype follicle cells and wildtype wing discs at wing-specific (green), follicle-specific (blue), and shared (black) transcripts (p < 0.0001; One way ANOVA). **F)** Histogram of replication timing log_2_ fold change of wing-specific (green) and follicle-specific (blue) transcripts.

To identify whether tissue-specific RT is driven by tissue-specific gene expression between wing discs and follicle cells, we directly compared differences in RT and gene expression at 10kb windows across the genome between the two tissues. First, we compared the average change in abundance of all transcripts within each window to the RT change of that window (Materials and Methods). Although transcript abundance was modestly elevated in wing discs versus follicle cells at windows of earlier RT in wing discs (average log_2_ fold change = 1.45CPM), we did not observe a strong correlation between elevated gene expression and earlier RT in follicle cells (Figure 2B,C; Figure S3B). These results were consistent whether we considered 1) the average change in the abundance of all transcripts overlapping each 10kb window (Figure 2B,C; Figure S3B), 2) the change of the most confident transcript (lowest p value) assigned to each window (Figure S3C), or 3) the change of the transcript with the greatest differential expression (absolute maximum log_2_ fold-change) assigned to each window (Figure S3D). Furthermore, 47.4% (791/1670) and 73.4% (813/1107) of windows with earlier RT in wing discs or follicle cells, respectively, do not contain a transcript with a significant increase in gene expression (Figure S3E), suggesting that tissue-specific RT and tissue-specific gene expression are mechanistically separable. Therefore, we conclude that differential gene expression between wing discs and follicle cells does not fully explain differences in RT between these two tissues.

As an independent method to assess the relationship between tissue-specific gene expression and RT, we identified genes expressed in both tissues (shared), genes expressed in wing discs only (wing-specific), and genes expressed in follicle cells only (follicle-specific) (Materials and Methods). We identified 12,626 genes that were expressed in both tissues, 901 genes that were wing-specific, and 517 that were follicle-specific (Figure 2D). When we quantified differential RT at both shared genes and tissue-specific genes, we observe earlier replication of wing-specific and shared genes in wing discs whereas follicle-specific genes do not replicate earlier in follicle cells (Figure 2E,F). These data again indicate that tissue-specific transcription and tissue-specific RT, although correlated, are separable. We hypothesized that earlier replication of shared genes in wing discs would correlate with elevated gene expression genome-wide in wing discs relative to follicle cells. Direct comparison of gene expression between the two tissues revealed a global increase of transcript abundance in wing discs relative to follicle cells (Figure S3F,G). Together, these data demonstrate that while gene expression and RT are correlated genome-wide (Figure S3H,I), changes in gene expression do not direct changes in RT between wing discs and follicle cells suggesting that RT and transcriptional activity are mechanistically separable.

### The mitotic-to endocycle transition does not affect DNA replication timing in follicle cells

The follicle cells of the adult ovary undergo a developmentally programmed cell cycle transition in which, after a series of mitotic divisions, they begin endocycling, a cell cycle consisting of S and G phases with no intervening mitoses (Figure 3A) (Edgar and Orr-Weaver 2001; Fox and Duronio 2013; Edgar et al. 2014). Follicle cells undergo three endocycles, resulting in a ploidy of 16C. Previous work has shown that there are distinct changes in genome regulation during the endocycle, including a global decrease in transcription, decrease in E2F1 target gene expression, and acquisition of endocycle-specific ORC binding sites (Maqbool et al. 2010; Sher et al. 2012; Hua et al. 2018; Rotelli et al. 2019). Therefore, we hypothesized that follicle cell replication timing may be influenced by this developmentally regulated cell cycle transition.

**Figure 3.**
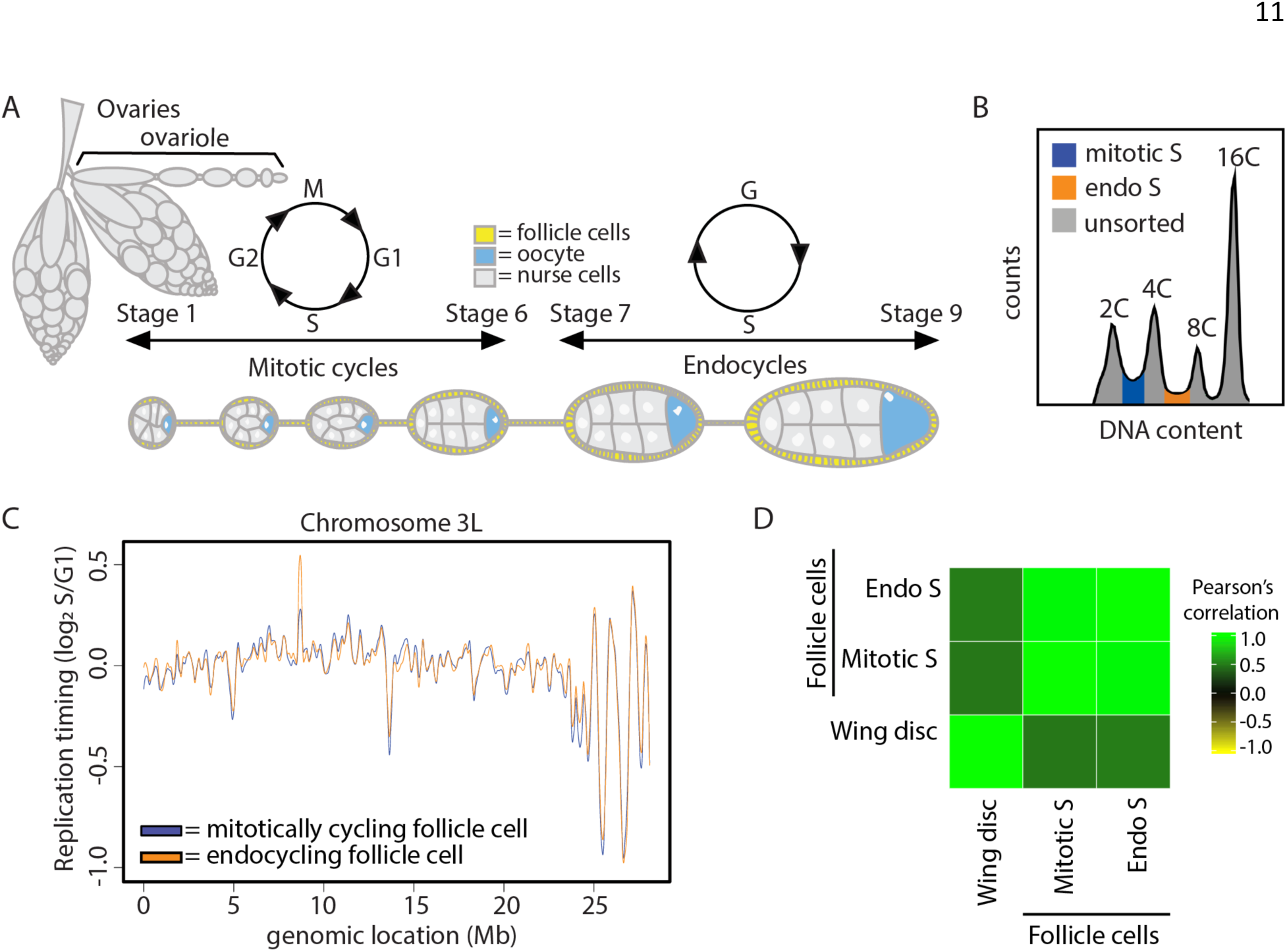
The mitotic to endocycle transition does not affect DNA replication timing within the follicle cells of the adult ovary. **A)** Early egg chamber development within the adult *Drosophila* ovary. **B)** Representative FACS profile of follicle cell nuclei isolated from whole ovaries. The 2C-4C S phase fraction (blue) are the mitotically cycling follicle cells, and the 4C-8C S phase fraction (orange) are the endocycling follicle cells. **C)** LOESS regression line showing average wildtype mitotically cycling follicle cells (blue) and wildtype endocycling follicle cells (orange) S/G1 (log_2_) replication timing values in at across the chromosome 3L scaffold. See Figure S4 for all other chromosome arms. **D)** Correlation matrix of S/G1 (log_2_) replication timing values for wildtype endocycling follicle cells (endo S), wildtype mitotically cycling follicle cells (mitotic S), and wild type wing discs.

To determine if the transition from a mitotic cycle to an endocycle causes a change in RT, we generated genome-wide replication timing profiles from wildtype endocycling follicle cells and compared them to the RT profiles we measured from wildtype mitotic follicle cells (Figure S4A,B). To this end, we collected the S phase populations between the 2C and 4C peaks (mitotic) and between the 4C and 8C peaks, which corresponds to the second of the three endocycles (Figure 3B). Direct comparison of RT profiles generated from wildtype mitotic (2C-4C) and endocycling (4C-8C) follicle cells showed no windows of differential RT genome-wide between the two populations of follicle cells (Figure 3C,D; Figure S4C; Table S1). Likewise, the gene expression profiles of these two populations of follicle cells were highly similar, with only six differentially expressed transcripts between mitotically cycling and endocycling follicle cells (p < 0.01, edgeR; Figure S2; Figure S4D). It is important to note that the first follicle cell endocycle likely initiates from G1 phase (Lilly and Spradling 1996; Calvi et al. 1998); therefore, the mitotic S phase sample may contain both mitotic and endocycling follicle cells. We were concerned that the impure cell population in the mitotic follicle cell dataset might mask any differential RT between the mitotic and endocycling populations. Based on the number of follicle cells in a mature egg chamber (~1000), we estimate that follicle cells in the first endo S phase could account for, at most, one half of the ‘mitotic’ follicle cell population (2C-4C) (Materials and Methods). Therefore, we performed an *in silico* false discovery rate (FDR) analysis by spiking in random reads from the wing disc RT dataset into the mitotic follicle cell RT dataset. Given that the endocycling follicle cells contribute no more than 50% of our total mitotic follicle cell population, we find that our analysis would be sensitive enough to accurately identify at least ~27% of the endocycle-specific RT differences (Figure S4E; Materials and Methods). Thus, endocycling S phase cells in the 2C-4C population do not mask a difference in RT between endocycling and mitotic follicle cells. Although we cannot exclude the possibility that minor changes in RT could be masked in in our data, we conclude that mitotic and endocycling follicle cells have remarkably similar RT profiles, arguing that cell lineage, not changes in the cell cycle, is a major contributing factor to RT.

### Rif1 fine tunes the replication timing program in different tissues

Rif1 is a global regulator of DNA RT from yeast to humans (Cornacchia et al. 2012; Hayano et al. 2012; Yamazaki et al. 2012; Peace et al. 2014; Seller and O’Farrell 2018). We sought to determine whether Rif1 regulates RT in a tissue-specific manner or whether Rif1-dependent RT domains are hardwired into the genome. To address these questions, we generated genome-wide RT profiles from mitotic follicle cells and wing discs in a *Rif1* null (*Rif1*^−^) mutant previously generated by our lab (Figure S5A,B; (Munden et al. 2018)). Individual replicates of *Rif1*^−^ RT data generated from either wing discs or follicle cells correlated well (Figure S5C; Figure S6A), whereas comparison of *Rif1*^−^ and wildtype RT data revealed that approximately 13% of the genome has differential RT in mitotically cycling follicle cells and 8% of the genome has differential RT in wing discs (Pearson’s correlation coefficient = 0.52 and 0.78, respectively; Figure S6B; Figure S5D). For the *Rif1*^−^ mutant follicle cells, 8.2% of windows displayed advanced RT while 5.0% of windows had delayed RT (Figure 4A-C; Figure S6C; Table S1). In the *Rif1*^−^ mutant wing disc, 4.1% of windows had advanced RT and 3.9% of windows had delayed RT (Figure 4A-C; Figure S5E; Table S1). Furthermore, the magnitude of RT changes within windows of differential RT between *Rif1*^−^ and wildtype was significantly greater in follicle cells than that observed in wing discs (Figure 4B,D). These data show that Rif1 has a greater impact on RT in follicle cells than wing discs, arguing that Rif1-dependent RT domains are not hardwired into the genome.

**Figure 4.**
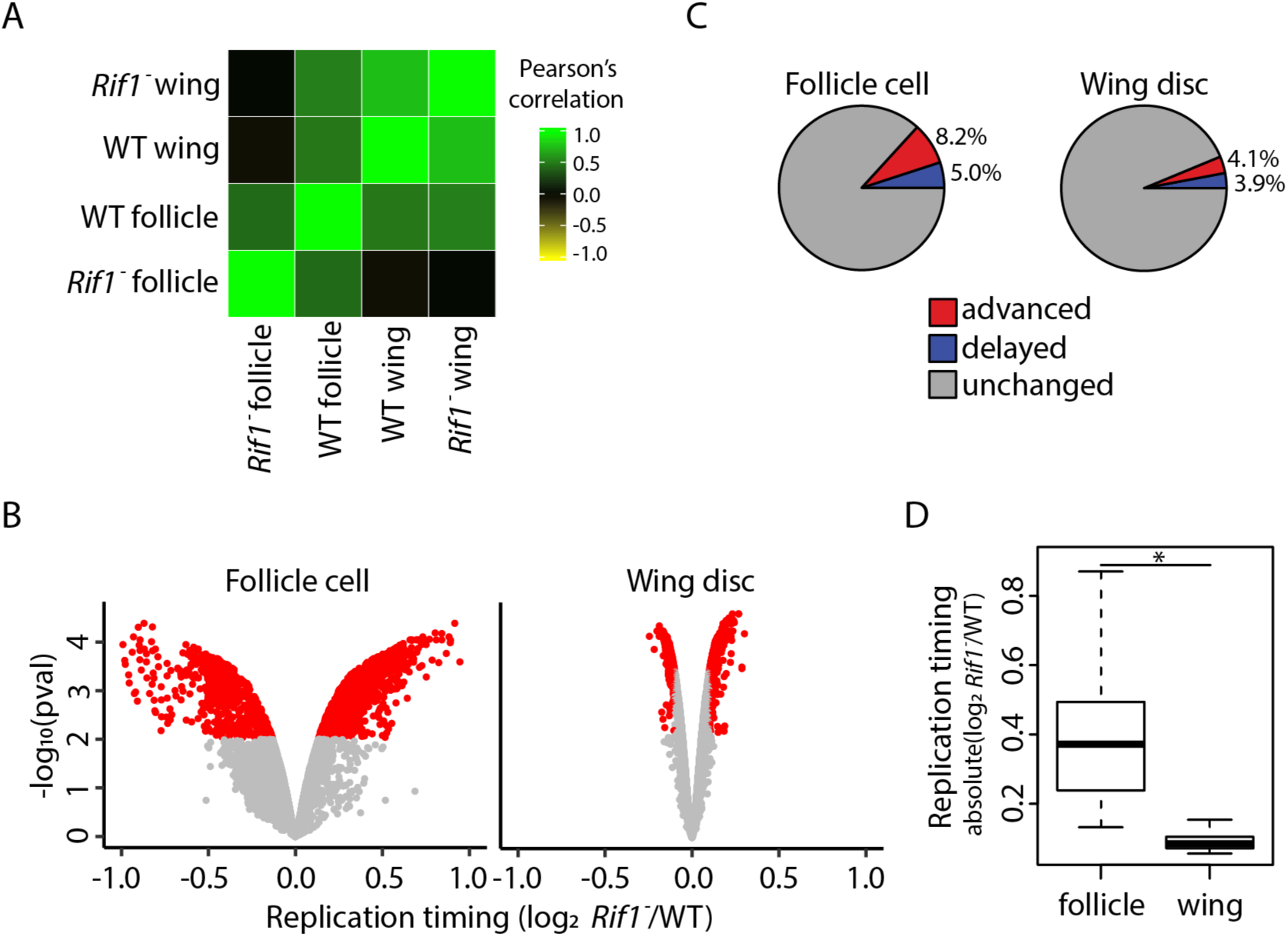
Rif1 regulates RT in a lineage-specific manner. **A)** Correlation matrix of S/G1 (log_2_) replication timing values for wildtype mitotically cycling follicle cells (WT follicle), *Rif1*^−^ mitotically cycling follicle cells (*Rif1*^−^ follicle), wildtype wing discs (WT wing), and *Rif1*^−^ wing discs (*Rif1*^−^ wing). **B)** Volcano plot of the *Rif1*^−^/control ratio of normalized replication timing values (S/G1 (log_2_)) plotted versus the -log_10_ p value (adjusted for multiple testing) in follicle cells (left) and wing discs (right). Significant replication timing changes are indicated (red; p < 0.01, absolute log_2_ fold change > 0.1; limma). **C)** Pie chart of all 100kb windows of significantly advanced RT (red), significantly delayed RT (blue), and unchanged RT (grey) across the major chromosome scaffolds in *Rif1*^−^ mutants relative to wildtype control in follicle cells (left) and wing discs (right) **D)** S/G1 (log_2_) absolute log_2_ fold change at 100kb windows of significant RT change between *Rif1*^−^ and control in follicle cells and wing discs (Student’s t test, p < 2.2 × 10^−16^).

Rif1 promotes late replication likely by preventing replicative helicase activation (Hayano et al. 2012; Davé et al. 2014; Hiraga et al. 2014; Mattarocci et al. 2014; Hiraga et al. 2017). Therefore, we hypothesized that advanced RT in a *Rif1*^−^ mutant is a direct effect of loss of Rif1 function, whereas delayed RT in a *Rif1*^−^ mutant is a secondary effect. This hypothesis predicts that when comparing different *Rif1*^−^ mutant cell types, there should be a greater extent of overlap between regions with advanced RT (direct) than between regions with delayed RT (indirect). We found that 43.8% (242/552) of windows with advanced RT in wing discs were also advanced in follicle cells. In contrast, only 16.9% (89/527) of windows with delayed RT in wing discs were also delayed in follicle cells (Figure 5A). These data support the hypothesis that advanced RT is a direct effect of Rif1 loss whereas delayed RT is likely a secondary effect.

**Figure 5.**
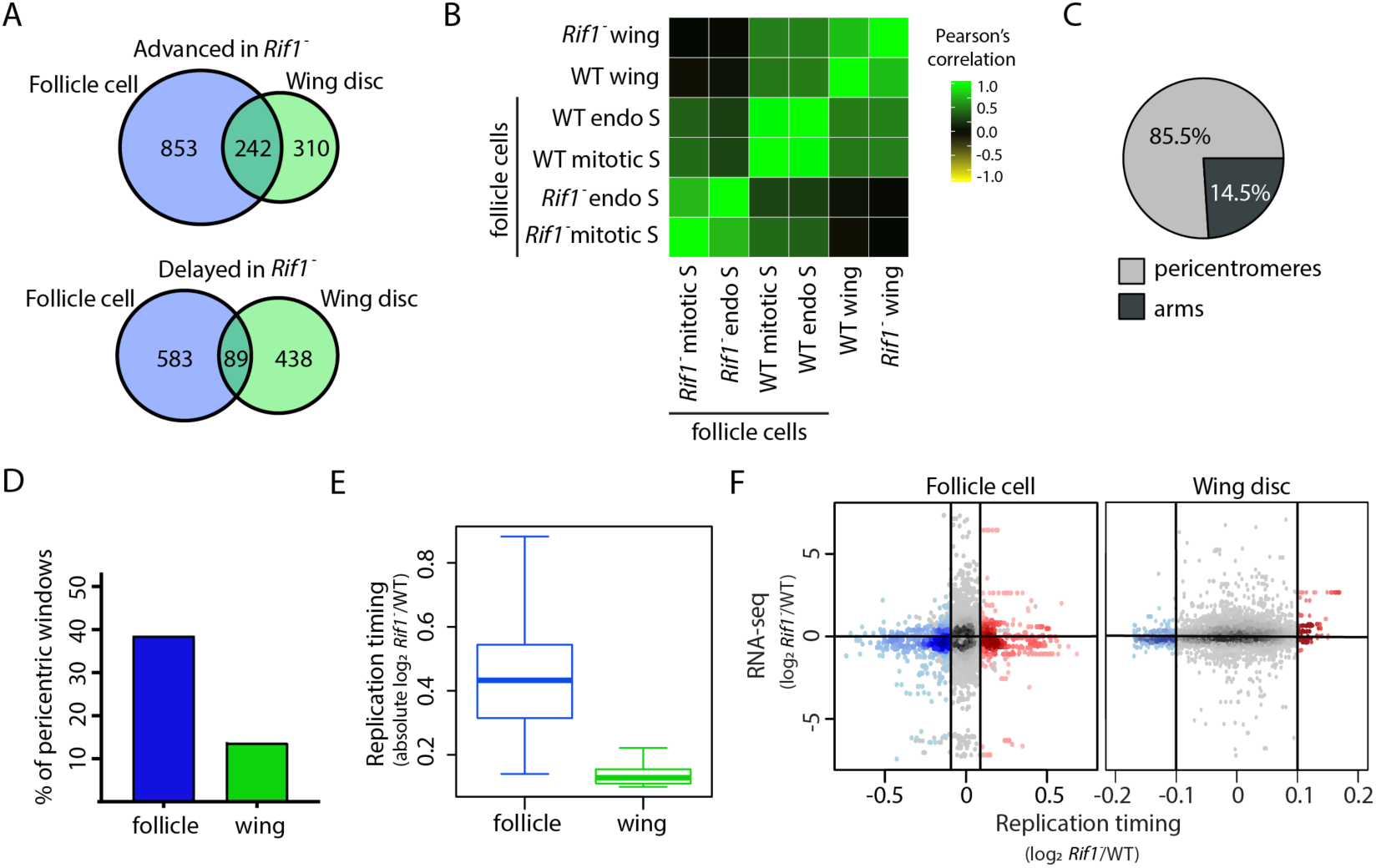
Rif1 promotes late replication of pericentric heterochromatin across lineages. **A)** Venn diagrams comparing significantly advanced (top) and delayed (bottom) 100kb windows identified in *Rif1*^−^ follicle cells (left; blue) and wing discs (right; green) (p<0.01 and absolute log_2_ fold change > 0.1; limma). **B)** Correlation matrix of S/G1 (log_2_) replication timing values for wildtype mitotically cycling follicle cells (WT mitotic S), *Rif1*^−^ mitotically cycling follicle cells (*Rif1*^−^ mitotic S), wildtype endocycling follicle cells (WT endo S), *Rif1*^−^ mitotically cycling follicle cells (*Rif1*^−^ endo S), wild type wing discs (WT wing), and *Rif1*^−^ wing discs (*Rif1*^−^ wing). **C)** Pie chart of all 100kb windows of commonly advanced RT between *Rif1*- wing discs and follicle cells. Windows within pericentromeres are in grey and chromosome arms are in black. **D)** Bar plot of the percentage of 100kb windows in pericentric heterochromatin with significantly advanced RT. **E)** S/G1 (log_2_) absolute log_2_ fold change at all 100kb windows located in pericentric heterochromatin between *Rif1*^−^ and control (Student’s t test, p < 2.2 × 10^−16^). **F)** Heatscatter plot of the *Rif1*^−^/control ratio of normalized replication timing values (S/G1 (log_2_)) plotted versus the *Rif1*^−^/control ratio of the most confident transcript (lowest p value) at each window across the major chromosome scaffolds. Significantly advanced (red) and delayed (blue) windows are indicated (p < 0.05, absolute log_2_ fold change > 0.1 (vertical lines); limma).

While measuring RT values for *Rif1*^−^ mutant and control samples, we profiled *Rif1*^*−/+*^ heterozygous follicle cells (Figure S7A,B). To our surprise, this heterozygous genotype displayed an intermediate RT phenotype with 3.6% (478/13391) of windows with advanced RT and 1.6% of windows with delayed RT relative to wildtype follicle cells (Figure S7C). Furthermore, 87.0% of windows with significantly advanced and 57.5% with significantly delayed RT in *Rif1*^−^ heterozygotes were also affected in *Rif1*^−^ follicle cells, indicating dependency on Rif1 function (Figure S7D). These data demonstrate that *Rif1* is haploinsufficient for RT control.

As an independent metric to address the specificity of commonly advanced and/or delayed RT changes, we asked whether common RT changes between mitotic follicle cells and wing discs were also detected in *Rif1*^−^ endocycling follicle cells. We generated RT profiles from *Rif1*^−^ endocycling follicle cells and found that individual replicates of RT data correlated well (Figure S8A). In contrast, 14.8% of windows displayed differential RT in *Rif1*^−^ endocycling follicle cells relative to control with 7.2% being advanced and 7.6% being delayed (Figure 5B; Figure S8B; Table S1). Although RT was similar between wildtype mitotic and endocycling follicles cells, a *Rif1* mutation affected these cell populations differently. We found that 72.1% (789/960) of advanced windows in *Rif1*^−^ endocycling follicle cells were also advanced in *Rif1*^−^ mitotic follicle cells, and only 37.9% (388/1024) of the windows that were delayed in *Rif1*^−^ endocycling follicle cells were also delayed in *Rif1*^−^ mitotic follicle cells (Figure S8C). Accordingly, the low degree of overlap between windows of delayed RT is reflected by the low genome-wide RT correlation between *Rif1*^−^ mitotic and endocycling follicle cells (Figure 5B; Figure S8D). Interestingly, many of the regions of advanced RT changes that were in common between *Rif1*^−^ wing discs and mitotic follicle cells were also detected in *Rif1*^−^ endocycling follicle cells while the delayed RT changes were mostly non-overlapping (72.7% (176/242) and 47.2% (42/89), respectively). Therefore, while Rif1 regulates RT in a tissue-specific manner, Rif1 appears to regulate RT in a core region of the genome regardless of cell type.

### Rif1 controls RT of pericentric heterochromatin

Almost all commonly advanced windows in *Rif1*^−^ mutant cell populations are located within pericentric heterochromatin, where Rif1 is known to localize (Buonomo et al. 2009; Munden et al. 2018; Seller and O’Farrell 2018). In contrast, all but eight of the commonly delayed windows are located along euchromatic chromosome arms (Figure 5C; Figure S9A). This relationship is also true for tissue-specific RT changes in *Rif1*^−^ wing discs and follicle cells—advancements are over-represented in pericentric heterochromatin whereas delays are over-represented along chromosome arms (Figure S9B). Collectively, these data suggest that Rif1 directly regulates late replication and may play a significant role in regulating late replication of pericentric heterochromatin. Interestingly, almost 40% of pericentric heterochromatin advances in *Rif1*^−^ follicle cells (both mitotically cycling and endocycling), whereas 2.8-fold fewer pericentric windows advance RT in *Rif1*^−^ wing discs (Figure 5D; Figure S9B). Furthermore, the overall RT of *Rif1*^−^ pericentric heterochromatin remains very late in wing discs relative to the average RT of the chromosome arms, and the magnitude of RT advancement is less than that observed in *Rif1*^−^ pericentric heterochromatin in follicle cells (Figure 5E; Figure S5E). Therefore, Rif1 contributes more substantially to late replication of pericentric heterochromatin in follicle cells than in wing discs.

Some genomic regions of *Drosophila* endocycling cells are under-replicated relative to the rest of the genome; i.e. they have reduced copy number relative to overall ploidy. This is particularly true in pericentric heterochromatin in salivary glands, and this under-replication requires Rif1 (Munden et al. 2018). Consequently, because our RT protocol measures relative copy number in S phase versus G1 phase, one possible explanation for the significantly earlier replication of pericentric heterochromatin in polyploid *Rif1*^−^ follicle cells relative to diploid *Rif1*^−^ wing discs is a loss of under-replication of pericentric heterochromatin. Multiple observations, however, indicate that we are measuring true changes in RT rather than the loss of under-replication in *Rif1*^−^ follicle cells. First, loss of under-replication predicts that 100% of pericentric heterochromatin would be scored as “advanced” RT. However, we found that only 40% of pericentric heterochromatin advances RT in *Rif1*^−^ mitotic and endocycling follicle cells (Figure 5D; Figure S8B). Second, if pericentric heterochromatin was under-replicated in wild type endocycling follicle cells, we would expect to observe a reduced copy number in pericentric heterochromatin relative to wildtype mitotically cycling follicle cells. However, pericentric heterochromatin copy number profiles derived from wildtype mitotic and endocycling S phase fractions are not different from one another (Figure S10). Together, these data support the conclusion that Rif1 regulates RT uniquely in different cell types and that the RT differences measured in *Rif1*^−^ follicle cells represent changes in RT and do not result from changes in under-replication.

### Rif1 controls RT independently of gene expression

To determine whether RT changes in *Rif1*^−^ wing discs and follicle cells were due to transcriptional deregulation, we generated transcriptomes from *Rif1*^−^ follicle cells and *Rif1*^−^ wing discs. We identified only 121 and 60 differentially expressed transcripts between *Rif1*^−^ and controls in wing discs and mitotic follicle cells, respectively, demonstrating that gene expression is largely unaffected after loss of Rif1 function (Figure S6D). We found only 2.1% (28/1342) of differential RT windows in follicle cells and 19.5% (99/507) of differential RT windows in wing discs contain at least one differentially expressed transcript (Figure 5F). Together, these data show that while loss of Rif1 function affects RT to a greater extent in follicle cells relative to wing discs, these RT changes likely do not result from transcriptional deregulation.

### Rif1’s PP1 binding motif is essential for Rif1-mediated RT control

Rif1 impacts the RT of pericentric heterochromatin to a greater extent in follicle cells than in wing discs (Figure 5D,E), suggesting a different requirement for Rif1 in RT regulation of pericentric heterochromatin in different tissues. To further understand these mechanistic differences, we assessed what role the PP1 binding motif within Rif1 has on RT control of pericentric heterochromatin in wing discs and follicle cells. Rif1 orthologs from yeasts to humans contain a PP1 binding motif, and mutation of this motif prevents Rif1 association with PP1 in multiple systems ((Davé et al. 2014; Hiraga et al. 2014; Mattarocci et al. 2014; Sreesankar et al. 2015; Alver et al. 2017; Hiraga et al. 2017; Sukackaite et al. 2017)). We previously generated an allele of Rif1 (*Rif1*^*PP1*^) where the conserved SILK/RSVF PP1 interaction motif is mutated to SAAK/RASA (Munden et al. 2018). We generated genome-wide RT profiles from *Rif1*^*PP1*^ wing discs and follicle cells. Individual replicates from each tissue correlated well (Pearson’s correlation = 0.91 and 0.89; Figure S11A,B; Figure S12A,B). In contrast, we found that 17.9% and 11% of windows in *Rif1*^*PP1*^ wing discs and follicle cells, respectively, displayed differential RT relative to control (Figure 6A,B; Figure S11C,D; Figure S12C,D; Table S1). Strikingly, *Rif1*^*PP1*^ wing discs displayed over 3-fold the number of advanced windows compared to *Rif1*^−^ wing discs. In addition, almost all (94.4%) advanced windows in *Rif1*^−^ wing discs were also advanced in *Rif1*^*PP1*^ mutants (Figure 6B). Interestingly, in follicle cells, there was almost a complete overlap of advanced RT windows between *Rif1*^*PP1*^ and *Rif1*^−^ mutants. These data suggest that the *Rif1*^*PP1*^ and *Rif1*^−^ mutations potentially affect RT through different mechanisms in wing discs and through the same mechanism in follicle cells. In contrast, the overlap of delayed RT changes between *Rif1*^*PP1*^ and *Rif1*^−^ wing discs or follicle cells is poor (Figure 6B). These data further support that advanced RT in *Rif1* mutants is a direct consequence of Rif1 loss, whereas delayed RT is likely secondary effect.

**Figure 6.**
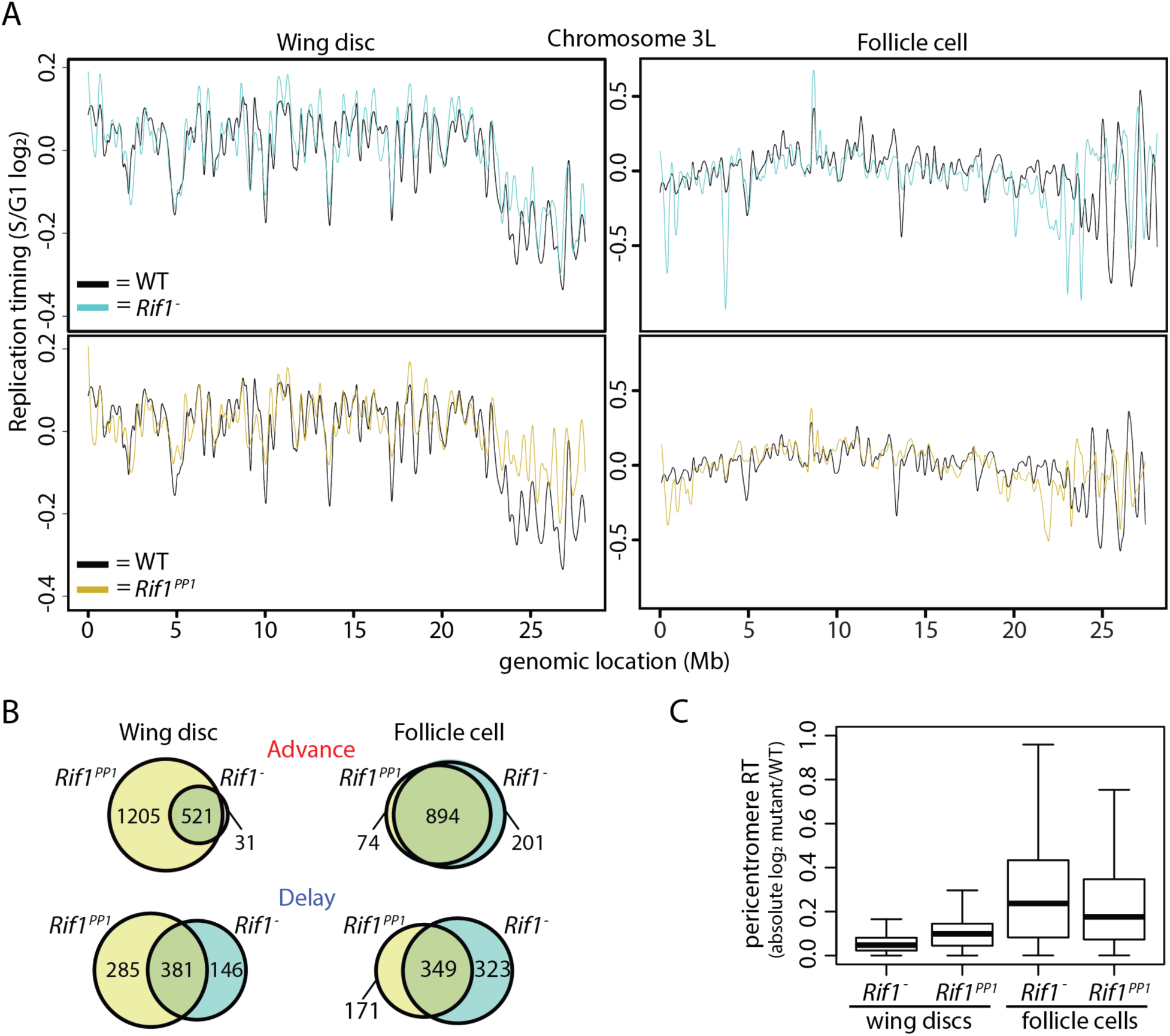
Rif1’s PP1 binding motif is essential for Rif1-mediated RT control. **A)** LOESS regression line showing average *Rif1*^−^ (cyan), *Rif1*^*PP1*^ (gold), and wildtype (black) S/G1 (log_2_) replication timing values in wing discs (left) and follicle cells (right) across the chromosome 3L scaffold. See Figures S5, S6, S11, and S12 for other chromosomes. **B)** Venn diagrams comparing significantly advanced (top) and delayed (bottom) 100kb windows identified in *Rif1*^−^ (cyan) and *Rif1*^*PP1*^ (gold) wing discs (left) and follicle cells (right) (p<0.01 and absolute log_2_ fold change > 0.1; limma). **C)** Box plot of absolute mutant/control log_2_ ratio of normalized replication timing values (S/G1 (log_2_)) at all pericentromeric regions of the major chromosome scaffolds.

As Rif1 affects RT of pericentric heterochromatin in both tissues, we hypothesized that RT changes in *Rif1*^*PP1*^ tissues would preferentially be located at pericentromeres. We found that approximately 48% of pericentric heterochromatin displayed a significant advancement of RT in *Rif1*^*PP1*^ wing discs, unlike what we found for *Rif1*^−^ null wing discs where only ~10% of pericentric heterochromatin advanced. The *Rif1*^*PP1*^ wing disc RT phenotype is more similar to what we observed at pericentric heterochromatin in *Rif1*^−^ follicle cells (Figure 5A). Specifically, 80% (876/1095) of advanced windows in *Rif1*^−^ mitotic follicle cells were also advanced in *Rif1*^*PP1*^ wing discs (Figure S12E). Additionally, all commonly advanced windows between *Rif1*^−^ follicle cells and wing discs were advanced in *Rif1*^*PP1*^ wing discs. Interestingly, while the magnitude of RT change at pericentromeres is significantly greater in *Rif1*^*PP1*^ wing discs relative to *Rif1*^−^ wing discs (p < 2.2 × 10^−16^), the magnitude of RT change in *Rif1*^*PP1*^ wing discs remains significantly lower than what is observed in *Rif1*^−^ or *Rif1*^*PP1*^ follicle cells (Figure 6C). Collectively, these data demonstrate that the *Rif1*^*PP1*^ mutation differentially affects pericentric heterochromatin RT relative to the *Rif1*^−^ mutation in wing discs and suggest that regulatory mechanisms, potentially including the Rif1-PP1 interaction, function differently to regulate late RT of pericentromeres between tissues.

## Discussion

Our findings provide insight into the relative contributions that cell type, gene expression, cell cycle, and Rif1 have on RT control. By comparing genome-wide RT profiles from unperturbed cells from distinct tissues, we demonstrated that cell lineage has a larger effect on RT than Rif1, an evolutionarily conserved regulator of RT. We also found that the RT program is not modified in response to the physiological and transcriptional changes that occur during the mitotic-to-endocycle transition and that transcriptional differences between cell types do not drive changes in RT.

We found that ~30% of the genome had different RT in the two tissue types we examined, and that transcriptional changes do not account for these changes. Studies in other systems also have failed to establish a direct relationship between changes in RT and changes in transcriptional activity (MacAlpine et al. 2004; Lubelsky et al. 2014; Siefert et al. 2017; Almeida et al. 2018; Armstrong et al. 2018). While transcriptional activity has long been correlated with RT, there are clearly mechanisms that control RT independently of transcription. RT is highly correlated with genome topology (Pope et al. 2014), and recent work has demonstrated that changes in TAD structure can be uncoupled from changes in gene expression (Ghavi-Helm et al. 2019). Therefore, our results are consistent with a model in which lineage-specific changes in genome topology, not transcription, underlie changes to the RT program as cells differentiate. These RT programs can then further be enforced by *trans*-acting factors such as Rif1.

When comparing different tissues, we found a higher degree of overlap between regions of the genome that transition from late-to-early in the absence of Rif1 than those that transition from early-to-late. These data imply that Rif1 directly promotes late replication of specific regions of the genome while indirectly affecting regions of the genome that normally replicate early. It is currently unknown, however, how Rif1 is targeted to heterochromatin and other late-replicating regions of the genome to delay RT. Rif1 dynamically associates with heterochromatin from yeasts to humans (Buonomo et al. 2009; Seller and O’Farrell 2018). In early *Drosophila* embryos, Rif1 is recruited to heterochromatic regions independently of HP1a, and then displaced from heterochromatin immediately before heterochromatin is replicated late in S phase (Seller and O’Farrell 2018). Chromatin immunoprecipitation of Rif1 followed by sequencing has revealed that in yeast and mouse cells Rif1 targets many other regions of the genome with both late and early replicating domains (Hayano et al. 2012; Foti et al. 2016). Our results argue that Rif1 localization to chromatin is likely influenced by cell type-specific factors.

Our results demonstrate that in metazoans the PP1 interaction motif of Rif1 can contribute to Rif1-mediated RT control. These data suggest that helicase inactivation, or inactivation of another PP1 target near origins of replication, is critical for Rif1-mediated RT control. Multiple models have been proposed to explain how Rif1 controls RT. First, through a direct interaction with PP1, Rif1 is thought to counteract DDK-mediated helicase activation and delay replication of Rif1-associated regions (Davé et al. 2014; Hiraga et al. 2014; Alver et al. 2017). Second, based on 4C experiments with five viewpoints, Rif1 was shown to affect chromatin contacts between different RT domains, suggesting that Rif1 controls RT through nuclear organization (Foti et al. 2016). It is unclear how these different models are related, if at all. Furthermore, while the timing decision point occurs in G1 phase, helicase activation occurs throughout S phase, raising additional mechanistic questions about how Rif1 controls RT. Recent work in budding yeast has shown that DDK can act in G1 phase (Zhang et al. 2019). Additionally, DDK-dependent helicase activation and Cdc45 recruitment in G1 phase is critical for the specification of certain replication origins. Thus, premature helicase activation in the absence of Rif1 during G1 phase could alter the localization of specific replication domains. While this model could unify the observations describing how Rif1 controls RT, further work is needed to test this possibility.

Our data suggest that different regulatory mechanisms control late RT between wing discs and follicle cells. The approximately 3-fold increase in the number of windows with advanced RT in *Rif1*^*PP1*^ wing discs relative to *Rif1*^−^ null wing discs was surprising. These data indicate that the presence of mutant Rif1^PP1^ protein results in a stronger effect than the absence of Rif1. One possibility is that Rif1^PP1^ acts in a dominant negative manner in regions of the genome that normally replicate late during S phase, such as pericentric heterochromatin. Another striking observation was that loss of Rif1 function in wing discs did not substantially advance RT in much of the pericentric heterochromatin. This result suggests that mechanisms in addition to Rif1/PP1-mediated MCM dephosphorylation act within the wing disc to promote late replication of pericentric heterochromatin.

In summary, our study demonstrates that cell lineage is a major driver of RT control within the context of a developing organism. Rif1 fine tunes the RT program established in different tissues, and each of these modes of RT control function independently of transcriptional control, suggesting additional levels of regulation.

## Materials and Methods

### FACS and genomic DNA sequencing

Isolated nuclei from *OregonR*, *Rif1*^*1*^/*Rif1*^*2*^ (*Rif1*^−^), and *Rif1*^*PP1*^/*Rif1*^*1*^ (*Rif1*^*PP1*^) female adult ovaries and *yw*, *Rif1*^−^, and *Rif1*^*PP1*^ female 3^rd^ instar larval wing imaginal discs from were sorted into G1 and S populations by a FACSAria II or III based on DAPI intensity and subsequently pelleted, flash frozen, and stored at −80°C prior to DNA isolation and library preparation. Libraries were prepared with the Rubicon ThruPLEX DNA-seq kit for wing imaginal disc samples and with the NEBNext Ultra II DNA Library Prep kit for follicle cell samples and subjected to Illumina HiSeq 2500 single-end 50bp sequencing for wing imaginal disc samples and Illumina HiSeq X or Novaseq 6000 paired-end 150bp sequencing for follicle cell samples.

### RT Characterization

Reads from G1 and S samples were aligned to the dm6 reference genome (Release 6.04) using Bowtie 2 (v2.3.2) default parameters (Langmead et al. 2009). Reads with a MAPQ score greater than 10 were retained using SAMtools (v1.9) (Li et al. 2009). BEDTools coverage (v2.26.0) was used to quantify the number of reads mapping to each 100kb window, with results normalized to read depth (Quinlan and Hall 2010). Replication timing (RT) values were obtained by averaging the S/G1 ratio of reads per million (RPM) value from each S phase replicate for a particular window size. Profiles were generated by plotting the RT value at each window versus genomic location. Quantile normalization was performed for comparisons between samples through the preprocess Core R package to equalize the dynamic range of RT values (Bolstad 2016). The limma statistical package was used to identify 100kb windows with significantly altered RT values (lmFit, p value adjusted for multiple testing (p<0.01); absolute log_2_ fold change > 0.1) (Newville et al. 2014). BEDTools intersect (v2.26.0) was used to determine overlap of 100kb windows with -f 0.5 and -u parameters (Quinlan and Hall 2010). RT values and limma-generated adjusted p values at 100kb windows were used to determine median RT values and adjusted p values at 10kb windows (BEDTools map v2.26.0), and the significance threshold was adjusted at 10kb windows (p value adjusted for multiple testing (p<0.05); absolute log_2_ fold change > 0.1) (Quinlan and Hall 2010). Coordinates of chromatin states were obtained from (Kharchenko et al. 2011) and converted to dm6 coordinates using the UCSC liftOver tool (Karolchik et al. 2004). To calculate RT domain sizes, we identified the genomic coordinates halfway between each peak and valley of an RT profile and determined the distance from one halfway point to the next.

For false discovery rate (FDR) calculations, spike-in RT bed files with 3 × 10^7^ reads were generated by combining either 3 × 10^5^ (1% impure), 1.5 × 10^6^ (5% impure), 3 × 10^6^ (10% impure), 7.5 × 10^6^ (25% impure), or 1.5 × 10^7^ (50% impure) randomly selected reads from each wing disc S phase replicate with 2.97 × 10^7^ (1% impure), 2.85 × 10^7^ (5% impure), 2.7 × 10^7^ (10% impure), 2.25 × 10^7^ (25% impure), or 1.5 × 10^7^ (50% impure) randomly selected reads from each mitotically cycling follicle cell S phase replicate. RT profiles generated from each test dataset (1% impure, 5% impure, 10% impure, 25% impure, and 50% impure) were directly compared to RT profiles from wing discs, and differential replication timing was identified as before using the limma statistical package (lmFit, p value adjusted for multiple testing (p<0.01); absolute log_2_ fold change > 0.1) (Newville et al. 2014). We estimate that 50% of the “mitotic” follicle cell population consists of endocycling follicle cells due to the following rationale: Because the total number of follicle cells in an egg chamber after the completion of the mitotic cell divisions is 1,024, the 2C-4C population used for sorting contains 2^10^ (1,024) mitotically cycling follicle cells from all egg chambers prior to Stage 7 per ovariole and (at most) 1,024 endocycling follicle cells from the Stage 7 egg chamber per ovariole.

### RNA Analyses

#### Follicle cell isolation, RNA extraction and sequencing

Follicle cells were isolated by trypsinizing ovaries from *OregonR* or *Rif1*^*1*^/*Rif1*^*2*^ females as described in (Cayirlioglu et al. 2003; Kim et al. 2011). Follicle cells were FACS sorted into TRIzol LS (Invitrogen) based on their ploidy and RNA was extracted according to the manufacture’s recommendation. 250,000 – 500,000 follicle cells were used per replicate. rRNA was depleted using the RiboMinus™ Eukaryote Kit for RNA-Seq (Invitrogen) and libraries were prepared using the NEBNext^®^ Ultra™ II RNA Library Prep.

#### Wing disc isolation, RNA extraction and sequencing

Total RNA was isolated from 40 *yw* and *Rif1*^*1*^/*Rif1*^*2*^ female 3^rd^ instar wing imaginal discs. Wing imaginal discs were homogenized in Trizol (Invitrogen) and flash frozen in liquid nitrogen. RNA was isolated using the Direct-zol RNA miniprep kit (Zymo Research). rRNA was depleted and libraries were prepared using the Ovation *Drosophila* RNA-Seq system (NuGEN). RNA isolated from *yw* wing imaginal discs was also made into libraries and sequenced with follicle cell RNA for all comparisons in Figure 2.

#### RNA seq analysis

TopHat default parameters (v2.1.1) (Trapnell et al. 2012) were used to align paired-end reads to the dm6 version of the *Drosophila* genome. Transcriptomes were generated using Cufflinks (v2.2.1, see supplementary materials for parameters). Differentially expressed transcripts were determined via edgeR statistical analysis (p value <0.01) (Robinson et al. 2010; McCarthy et al. 2012). For analyses comparison transcription to RT at 10kb windows, we either assigned the average RNA log_2_ fold change and average adjusted p-value from all transcripts overlapping each 10kb window or we assigned the log_2_ fold-change of the transcript with the lowest edgeR-generated p value at each 10kb window for analyses directly comparing RT and transcription. Results were similar irrespective of how transcription was assigned to RT windows.

### Data access

The data generated as a part of this study have been submitted to the NCBI Gene Expression Omnibus (GEO) under accession number GSE141632.

## Acknowledgements

This work was supported by NIH Grants R01-GM124201 to R.J.D and NSF MCB 1818019 to J.T.N. In addition, R.L.A. was supported in part by an NIH predoctoral training grant T32-GM007092. We thank the UNC Flow Cytometry and High Throughput Sequencing Core Facilities, supported in part by P30 CA016086 Cancer Center Core Support Grant to the UNC Lineberger Comprehensive Cancer Center. FACS results reported in this publication were supported in part by the North Carolina Biotechnology Center Institutional Support Grant 2012-IDG-1006. Flow Cytometry experiments were performed in the VMC Flow Cytometry Shared Resource. The VMC Flow Cytometry Shared Resource is supported by the Vanderbilt Ingram Cancer Center (P30 CA68485) and the Vanderbilt Digestive Disease Research Center (DK058404).

